# Activation of p21 limits acute lung injury and induces early senescence after acid aspiration and mechanical ventilation

**DOI:** 10.1101/2020.03.24.005983

**Authors:** Jorge Blázquez-Prieto, Covadonga Huidobro, Inés López-Alonso, Laura Amado-Rodriguez, Paula Martín-Vicente, Cecilia López-Martínez, Irene Crespo, Cristina Pantoja, Pablo J Fernandez-Marcos, Manuel Serrano, Jacob I Sznajder, Guillermo M Albaiceta

## Abstract

The p53/p21 pathway is activated in response to cell stress. However, its role in acute lung injury has not been elucidated. Acute lung injury is associated with disruption of the alveolo-capillary barrier leading to acute respiratory distress syndrome (ARDS). Mechanical ventilation may be necessary to support gas exchange in patients with ARDS, however, high positive airway pressures can cause regional overdistension of alveolar units and aggravate lung injury. Here, we report that acute lung injury and alveolar overstretching activate the p53/p21 pathway to maintain homeostasis and avoid massive cell apoptosis. A systematic pooling of transcriptomic data from animal models of lung injury demonstrates the enrichment of specific p53- and p21-dependent gene signatures and a validated senescence profile. In a clinically relevant, murine model of acid aspiration and mechanical ventilation, we observed changes in the nuclear envelope and the underlying chromatin, DNA damage and activation of the *Tp53*/p21 pathway. Absence of *Cdkn1a* decreased the senescent response, but worsened lung injury due to increased cell apoptosis. Conversely, treatment with lopinavir/ritonavir led to *Cdkn1a* overexpression and ameliorated cell apoptosis and lung injury. The activation of these mechanisms was associated with early markers of senescence, including expression of senescence-related genes and increases in senescence-associated heterochromatin foci in alveolar cells. Autopsy samples from lungs of patients with ARDS revealed increased senescence-associated heterochromatin foci. Collectively, these results suggest that acute lung injury activates p53/p21 as an anti-apoptotic mechanism to ameliorate damage, but with the side effect of induction of senescence.

## Introduction

The lungs have a stereotypic response to acute injury, which is preserved among species and many etiological agents. Once damage is inflicted, lung cells trigger a host response which can include inflammation, matrix remodeling and different forms of cell death, including apoptosis (Herold et al., 2013). Although a limited host response may help to clear the injurious agent and promote lung tissue repair (Blazquez-Prieto et al., 2018), an overexuberant host response can lead to severe injury and gas exchange worsening. Therefore, therapeutic strategies aimed to limit lung damage and interference with lung repair are important.

Lungs are exposed to mechanical load during every breath. In pathologic conditions, generation of higher pressure gradients necessary for adequate ventilation may cause excessive cell stretch (Perlman and Bhattacharya, 2007). This is especially relevant during mechanical ventilation with high pressures, which can lead to the so-called ventilator-induced lung injury (VILI) (Corbridge et al., 1990; Slutsky and Ranieri, 2013). In mechanically ventilated patients, a strategy aimed to limit VILI decreased mortality in patients with the acute respiratory distress syndrome (ARDS)(The Acute Respiratory Distress Syndrome Network, 2000).

Mechanotransduction is thought to regulate the molecular steps in VILI pathogenesis (Spieth et al., 2014). The nuclear envelope has been reported as an important cell mechanosensor and signal transducer (Swift et al., 2013). Mechanical stretch appears to increase Lamin-A in the nuclear envelope, leading to nuclear stiffening. These changes in the nuclear envelope can also activate p53-dependent pathways. Wildtype p53 is a master regulator of cell homeostasis and fate, and its activation may lead to a variety of responses, ranging from apoptosis to cell cycle arrest. Inhibition of this response has been shown to increase p21 (*Cdkn1a)* expression and decrease VILI in an experimental model (López-Alonso et al., 2018).

p53 and its downstream factor p21 are triggers of senescence (Varela et al., 2005), a cell response characterized by an stable arrest of the cell cycle and a switch towards a senescence-associated secretory phenotype (SASP). It has been proposed that senescence facilitates the clearance of damaged cells and is required for tissue repair (Muñoz-Espín and Serrano, 2014). Interestingly, some of the molecules have a significant overlap with the proinflammatory response associated with VILI.

We hypothesized that p53-dependent pathways play a role in the maintenance of lung homeostasis during acute injury, and that senescence could be a side effect of their activation. To test this hypothesis, we developed a clinically relevant model of lung injury caused by acid aspiration and VILI to assess the activation of p53 and its downstream factors.

## Results

### Transcriptomic signatures of p53/p21 activation and senescence in lung injury

To test the hypothesis that the p53/p21 pathway is activated during acute lung injury and to identify early markers of a switch towards a senescent phenotype, data from 11 datasets of mouse lung injury and mechanical ventilation (Supplementary Table 1) were pooled and gene expression analyzed (Figure 1A). Three different transcriptomic signatures related to p53-dependent upregulation (116 genes, 76 available in the pooled data), p21-dependent downregulation (14 genes, 8 available) (Fischer, 2017) and senescence (Hernandez-Segura et al., 2017) (55 genes, 44 available) were analyzed. A meta-score of expression of these genes was computed for each sample and compared to assess the effect of lung injury and mechanical ventilation.

**Figure 1.**
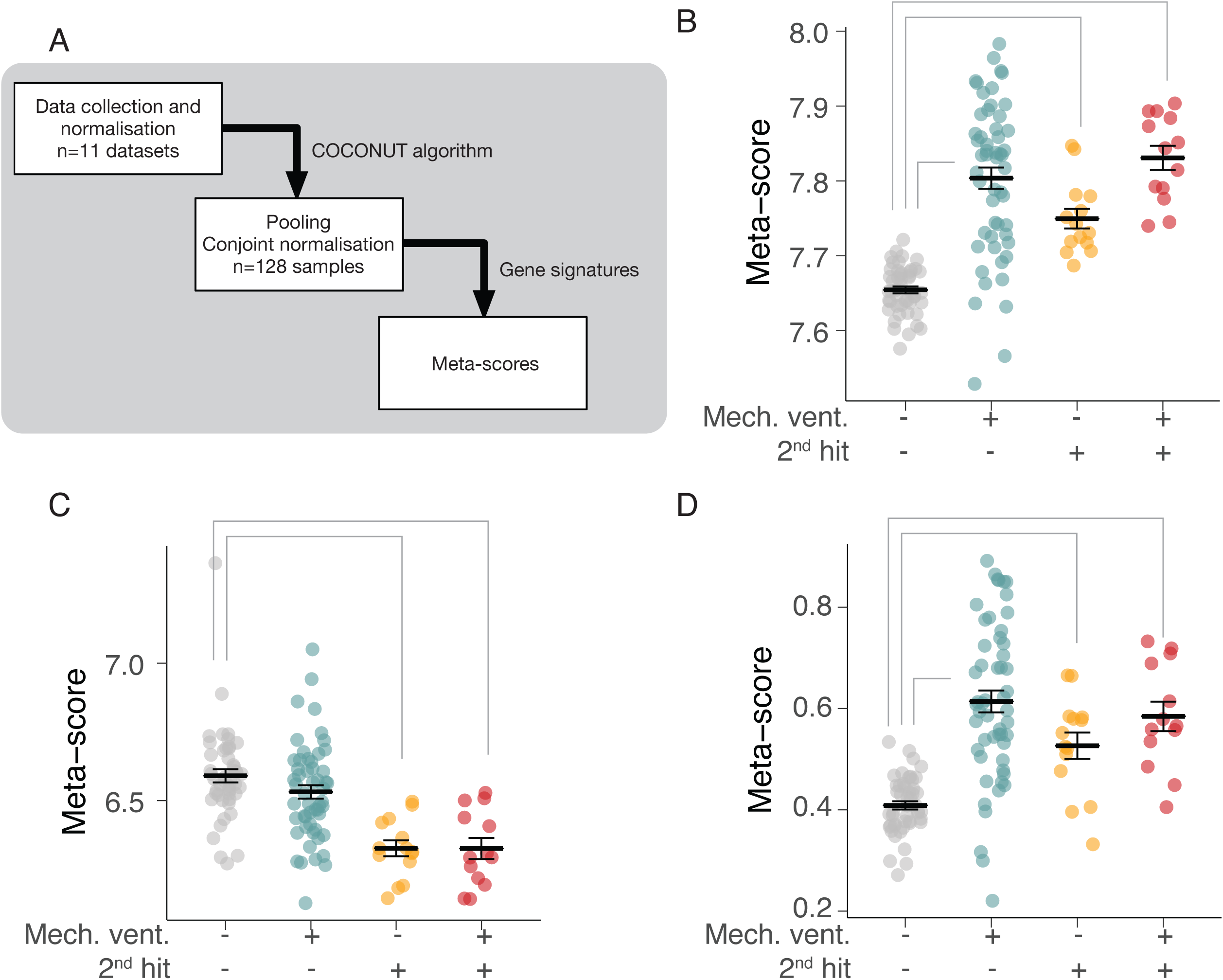
Expression of gene signatures. A: Overview of the analysis. Eleven datasets (128 samples) reporting gene expression in animal models of lung injury were pooled and analyzed to calculate different Meta-scores summarizing the expression of genes included in specific signatures. B: Meta-score of a p53-dependent signature for each experimental group (2^nd^ hit refers to any model of lung injury other than mechanical ventilation). C: Meta-score of a transcriptomic signature including genes downregulated by p21. D: Meta-score of a senescence-specific signature. Gray lines mark significant differences among groups (p<0.05 in Tukey’s post-hoc tests).

Animals subjected to acid aspiration lung injury and mechanical ventilation showed higher expression of p53-dependent genes (ANOVA p-value<0.001, Figure 1B), lower expression of p21-downregulated genes (ANOVA p-value<0.001, Figure 1C) and a higher metascore (Figure 1D) in the senescence signature than spontaneously breathing controls (ANOVA p-value<0.001, Figure 1D). The expression of each gene of these signatures is shown in Supplementary Figure 2. These results support the notion that lung injury and mechanical ventilation activate p53/p21 pathways and the molecular mechanisms of senescence in acutely injured lungs.

**Figure 2.**
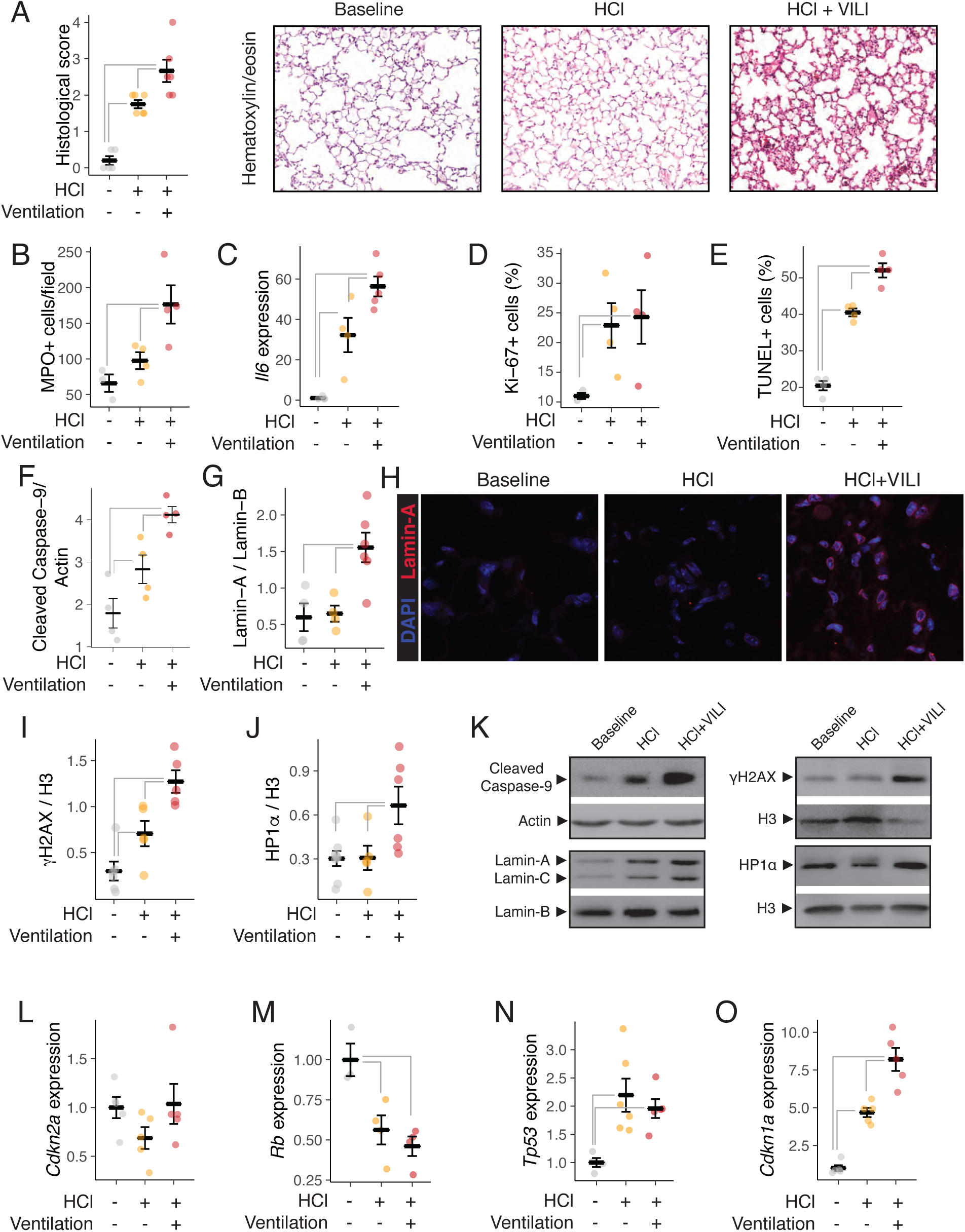
Characterization of lung injury. A: Acid instillation and mechanical ventilation caused lung damage assessed using a histological score. B: Myeloperoxidase-positive cell counts in histological sections, showing an increase of neutrophils in the injured lung. C: Expression of *Il6* in lung tissue. D: Quantification of Ki-67 positive cells in histological sections, as a marker of proliferation. E: TUNEL-positive cells in histological sections. F: Abundance of cleaved Caspase-9 in lung homogenates. G-H: Changes in Lamin-A/Lamin-B1 ratio in nuclei from lung tissue (G) and representative immunohistochemical sections (H). I-J: Abundance of γH2AX (I) and HP1α (J), markers of DNA damage and heterochromatin respectively, in nuclei from lung tissue. K: Representative western blots of the previous quantifications. L-O: Changes in expression of the canonical senescence inducers *Cdkn2a* (p16, L), *Rb* (M), *Tp53* (N) and *Cdkn1a* (p21, O). N=4-6 animals per group. Gray lines mark significant differences among groups (p<0.05 in Tukey’s post-hoc tests) E: Appearance of senescence-associated heterochromatin foci (SAHF), identified by their marker Macro-H2A, with lung injury and mechanical stretch. F-H: I: J: Gray lines mark significant differences among groups (p<0.05 in Tukey’s post-hoc tests)

### Activation of the p53/p21 pathway in a clinically relevant model

To explore the mechanisms involved in the activation of p53-dependent signals, an experimental model of acid aspiration- and mechanical ventilation induced lung injury was tested. Chlorhydric acid instillation and mechanical ventilation induced a significant increase in lung damage and inflammation, assessed by histological scores (Figure 2A), neutrophilic infiltrates (Figure 2B) and Il6 expression (Figure 2C). Lung damage increased the proportion of both proliferating and apoptotic cells in lung parenchyma (Figures 2D-E and Supplementary Figure 3). In line with this, the abundance of cleaved caspase-9 in lung tissue was increased with lung injury (Figure 2F)

**Figure 3.**
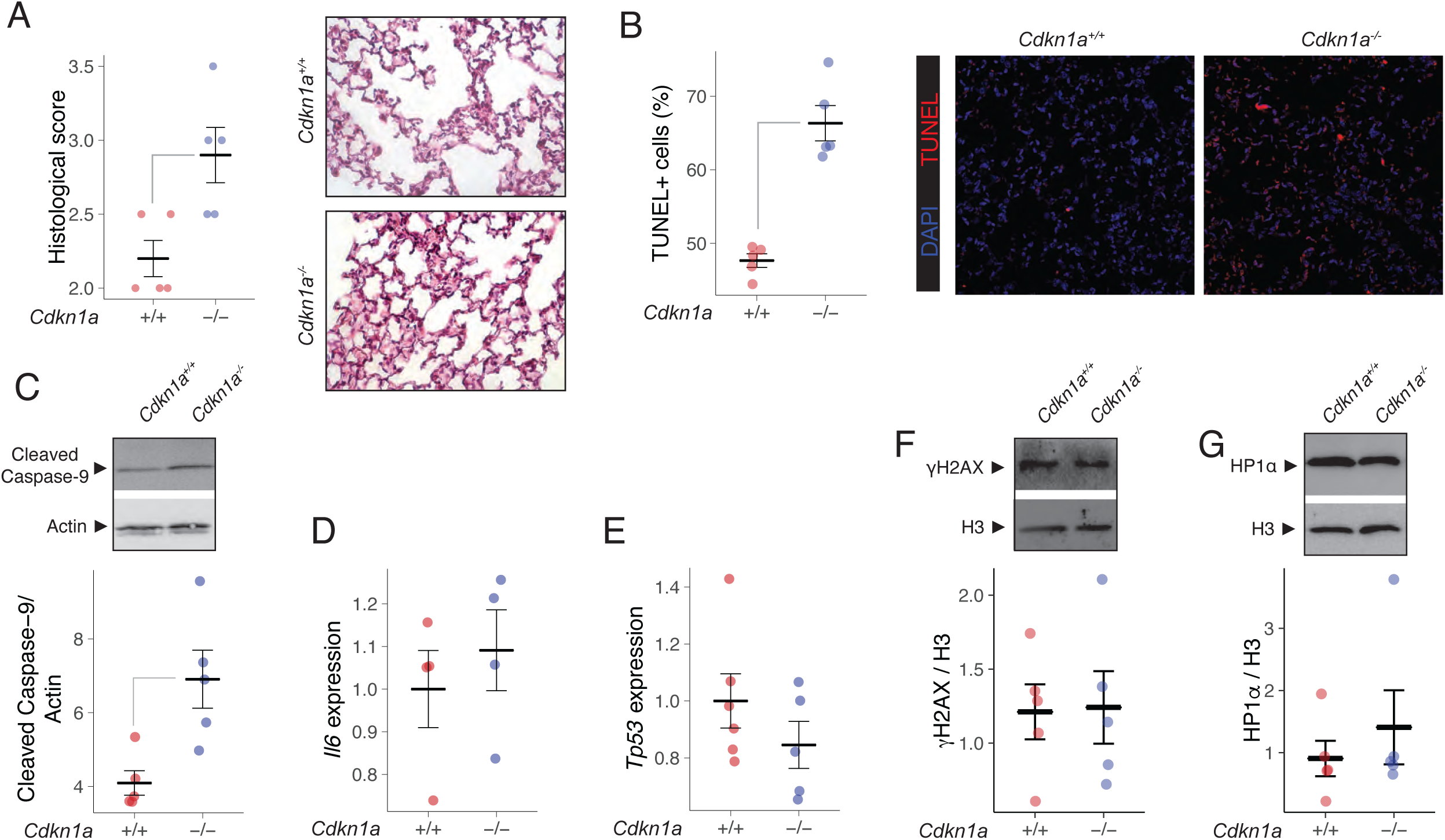
Lung injury in wildtype and *Cdkn1a*^-/-^ animals. A: Histological score of lung damage in both genotypes. B: Percentage of apoptotic (TUNEL+) cells. C: Abundance of cleaved caspase-9 in lung homogenates from both genotypes. D-E: Expression of Il6 (D) and *Tp53 (E)* in wildtype and mutant mice. F-G: Abundance of γH2AX (F) and HP1α (G), with representative western blots, in lung homogenates. N=4-6 animals per group. Gray lines mark significant differences among groups (p<0.05 in T tests).

We then explored the putative activators of this response to acute injury. Lamins in the nuclear envelope act as cell mechanosensors, regulating chromatin organization in response to mechanical stress. We observed that Lamin-A/Lamin-B ratio increased after mechanical stretch (Figure 2G). Immunofluorescence studies confirmed the increase in Lamin-A in the nuclear envelope after mechanical stretch, but not after hydrochloric acid instillation alone (Figure 2H). These changes in the nuclear envelope coexisted with an increase in γH2AX (Figure 2I) and HP1α (Figure 2J), markers of DNA damage and chromatin remodeling respectively, in nuclear extracts from ventilated animals. Panel 2K shows representative Western blots of these parameters.

The expression of the canonical responders to DNA damage *Cdkn2a* (p16) and *Tp53* (p53) and their corresponding downstream factors *Rb* and *Cdkn1a* (p21) was also assessed. There were no differences in the levels of *Cdkn2a* (Figure 2L), whereas expression of *Rb* was significantly decreased (Figure 2M). However, we observed significant increases in *Tp53* (Figure 2N) and *Cdkn1a* (Figure 2O) expression with lung injury.

### Increased lung damage in mice lacking p21

To address the role of p53 and p21 in acute lung damage, *Tp53*^-/-^, *Cdkn1a*^-/-^ mice and their wildtype counterparts were subjected to acid instillation followed by mechanical ventilation. In preliminary experiments, absence of *Tp53* did not modify lung injury (histological score 2.4±1.6 vs 2.3±1.2, n=4/group, p=0.91, Supplementary Figure 3), so we focused on the downstream factor p21. In absence of p21, the mice had worse lung injury (Figure 3A) and higher counts of apoptotic cells (Figure 3B) and cleaved caspase-9 abundance (Figure 3C) compared to their wildtype counterparts. There were no differences in *Il6* or *Tp53* expression nor in abundance of γH2AX or HP1α between genotypes (Figures 3D-G respectively).

### Lopinavir increases p21 and decreases lung damage

We have previously shown that HIV-protease inhibitors modify the nuclear response to mechanical stretch and protect against VILI (López-Alonso et al., 2018), an effect that could be due to the inhibition of the Lamin-A protease ZMPSTE24 (Coffinier et al., 2007). In our double-hit model, treatment with lopinavir/ritonavir impaired the structure of the nuclear lamina, decreasing the abundance of Lamin-A (Figure 4A), and decreased lung injury (Figure 4B), apoptotic cell count (Figure 4C) and cleaved caspase-9 (Figure 4D). Although abundance of γH2Ax was not modified by this treatment (Figure 4E), there was a marked decrease in HP1α (Figure 4F). Panel 4G shows representative blots of these measurements. Finally, treatment with lopinavir/ritonavir caused an increase in Il6 expression (Figure 4H), with no changes in *Tp53* expression (Figure 4I) but an increase in *Cdkn1a* (p21, Figure 4J).

**Figure 4.**
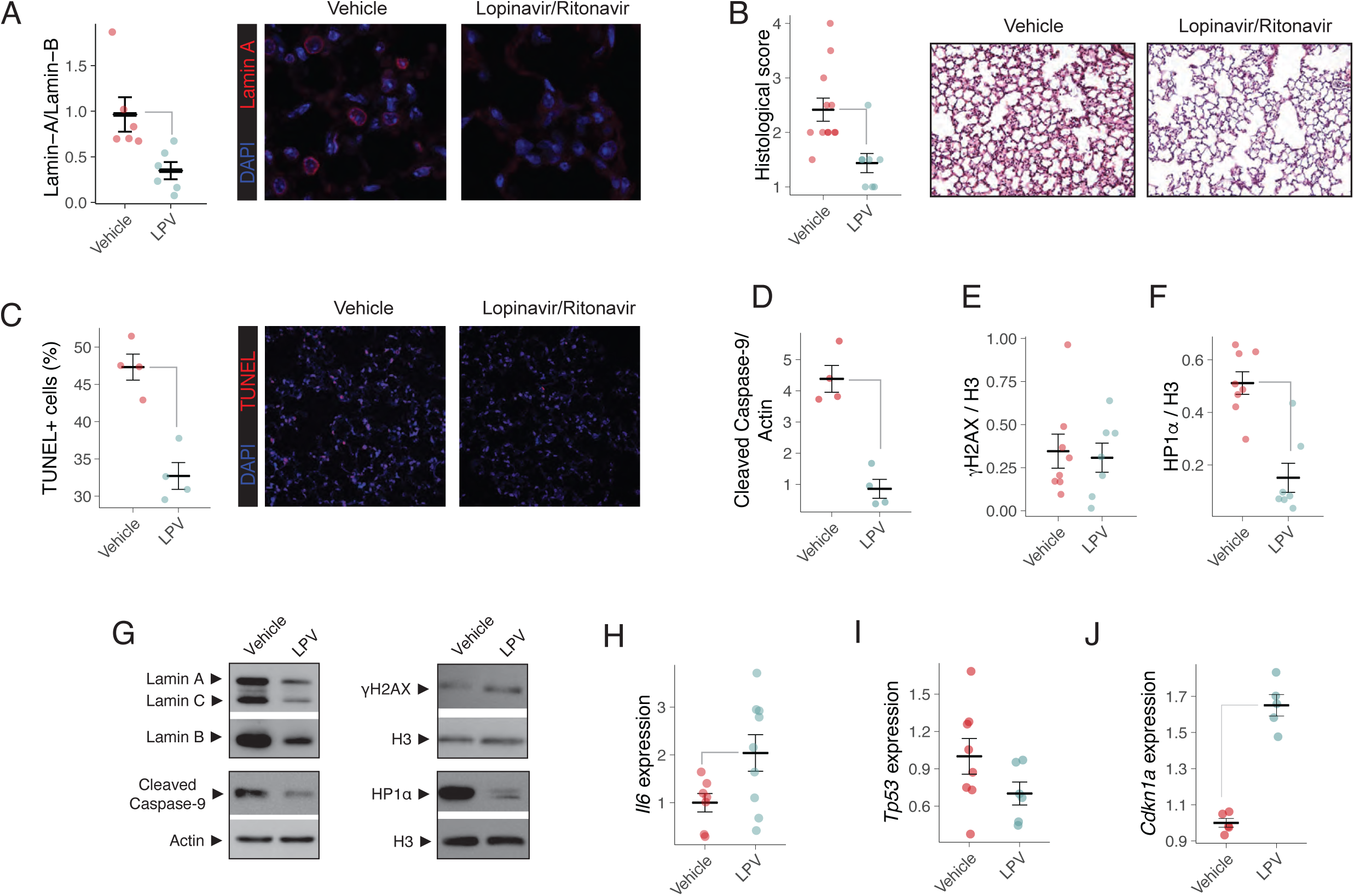
Effects of Lopinavir/Ritonavir on lung injury. A: Lamin-A abundance and staining in vehicle- and lopinavir/ritonavir treated animals. B: Histological score of lung damage. C: Apoptotic (TUNEL+) cell counts in both groups. D-F: Abundance of Caspase-9 in tissue homogenates (D), γH2AX (E) and HP1α (F), with representative western blots (G) in lung homogenates. H-J: Expression of Il6 (H), *Tp53* (I) and *Cdkn1a* (p21, J). N=7-10 animals per group.Gray lines mark significant differences among groups (p<0.05 in T tests).

### Early markers of senescence in acute lung injury

Then, we tried to identify early markers of senescence in our acute model. Acid aspiration and mechanical ventilation-induced lung injury was associated with an increase in the number of nuclei positive for Macro-H2A, a marker of senescence-associated heterochromatin foci (SAHF, Figure 5A), and changes in *Plk3* (Figure 5B), *Gdnf* (Figure 5C), and *Meis1* (Figure 5D), the genes from the senescence signature with the highest differential expression in the previous pooled analysis. These markers of senescence were modified by manipulation of the p21 pathway. Mutant animals lacking *Cdkn1a* exhibited a decreased number of SAHF after acid instillation and mechanical ventilation (Figure 5E) and in expression of the senescence-related gene *Plk3* (Figure 5F). In opposite, treatment with lopinavir/ritonavir (that increased *Cdkn1a* expression, Figure 4J) was related to lower counts of SAHF (Figure 5G), but increased *Plk3* expression (Figure 5H) Finally, to confirm the incidence of SAHF in patients, lung tissue from autopsies of critically-ill patients with and without lung injury and mechanical ventilation (Supplementary Table 2) were stained with antibodies against Macro-H2A. There were no differences in age between the three groups of patients (62±6, 61±11 and 54±10 years for patients without ARDS or mechanical ventilation, with ARDS but without mechanical ventilation and ARDS and ventilation respectively, p=0.17 in ANOVA). Similarly to the animal model, nuclear Macro-H2A increased in those with severe lung injury and mechanical ventilation (Figure 5I).

**Figure 5.**
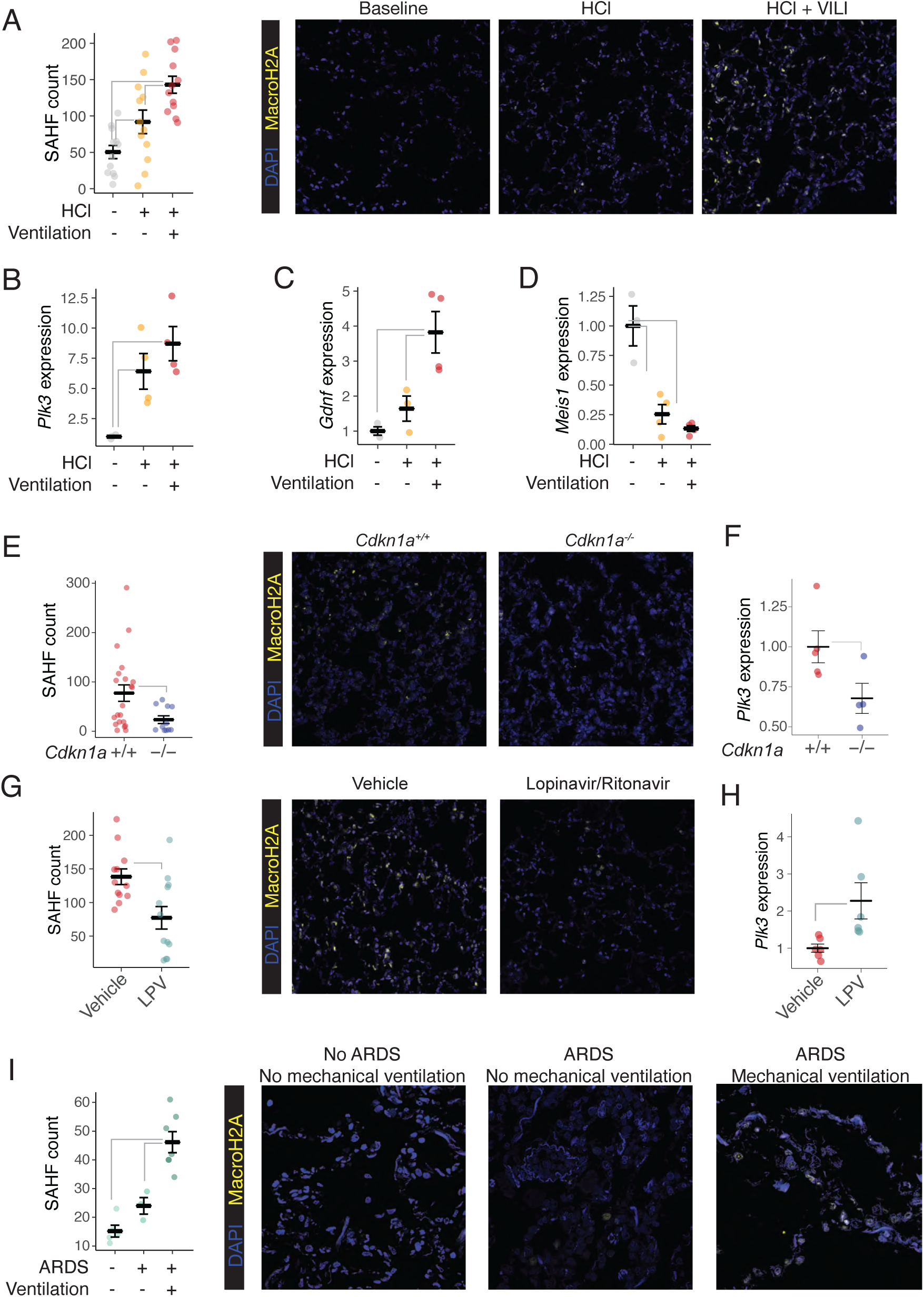
Identification of early senescence markers in experimental models and patients. A: Counts of Senescence-associated heterochromatin foci (SAHF) in the experimental model of lung injury of acid instillation and mechanical ventilation. B-D: Expression of *Plk3, Gdnf*, and *Meis1* in lung tissue. These senescence-associated genes were identified in the genomic analysis as those with the largest differences between control and injured samples. E: SAHF counts (E) and *Plk3* expression (F) in wildtype and *Cdkn1a*^-/-^ mice after lung injury. G-H: SAHF counts (G) and Plk3 expression (H) in vehicle and lopinavir/ritonavir (LPV)-treated mice after lung. I: Appearance of SAHF in autopsy samples from critically ill patients who died in the Intensive Care Unit with or without mechanical ventilation and acute respiratory distress syndrome (ARDS). N=4-7 animals per group, with three slides per animal as technical replicates in SAHF counts. Gray lines mark significant differences among groups (p<0.05 in Tukey’s post-hoc or in T tests).

Collectively, these findings suggest that lung injury and mechanical stretch trigger the appearance of early markers of senescence. The severity of lung injury and the abundance of senescence markers showed an inverse correlation after manipulation of p21 levels.

## Discussion

We provide evidence that acute lung injury and its treatment with mechanical ventilation alters the nuclear envelope and causes DNA damage, activating the p53/p21 pathway. Activation of p21 plays a homeostatic role, limiting the extent of apoptosis in response to injury. Moreover, this effect can be pharmacologically activated to ameliorate lung injury in a clinical setting. In spite of this beneficial effect, this pathway also leads to the appearance of early markers of senescence in lung tissue. Figure 6 summarizes the findings of this work.

**Figure 6.**
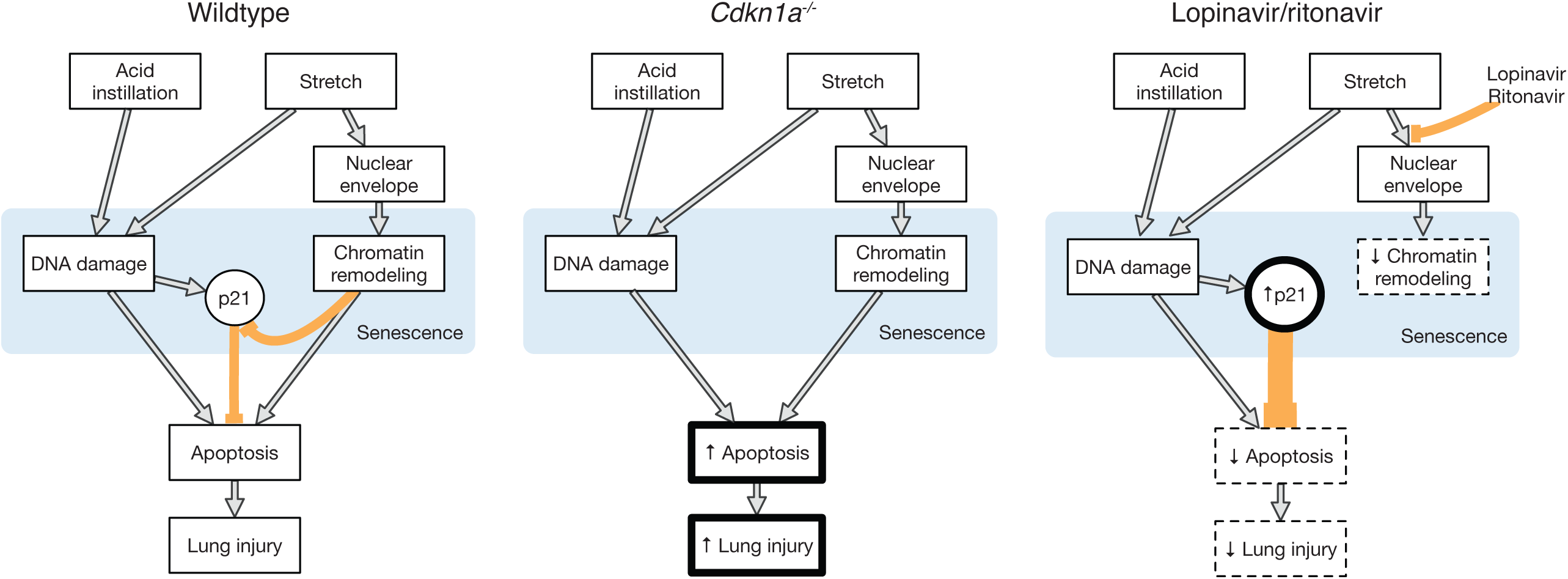
The role of p21 pathway on apoptosis and senescence after acute lung injury. A: In control mice, lung injury and mechanical stretch cause DNA damage and changes in the nuclear envelope, activating the cell senescence program. The amount of apoptotic cells depends on the equilibrium between the activation of pro-apoptotic responses triggered by injury itself and the anti-apoptotic effects of the senescence inducer *Cdkn1a* (p21). B: In mice lacking *Cdkn1a*, absence of this anti-apoptotic factor leads to an increase in apotosis and a more severe lung injury. C: Treatment with Lopinavir/ritonavir blocks the Lamin-A mediated chromatin remodeling, triggering a senescence-like response that increases p21 expression, thus decreasing apoptosis and lung damage.

### The p53/p21 axis in acute injury

P53 and its downstream transcription factor p21 are major regulators of cell homeostasis. It has been shown that p53 regulates permeability in lung endothelial cells after an inflammatory insult (Barabutis et al., 2018). Similarly, activation of this pathway in response to hypertonic saline decreased lung injury and inflammation in human airway epithelial cells (Gamboni et al., 2016). However, our observations in *Tp53*^-/-^ mice showed no differences in lung injury. Given the pleiotropic effects of p53 in cell homeostasis, this could be due to the existence of both protective and pathogenetic mechanisms.

One of the main effects of the cyclin kinase inhibitor p21 is the blockade of apoptosis (Fielder et al., 2017). It has been proposed that caspase-9 is the downstream target responsible for the anti-apoptotic effects of p21 (Sohn et al., 2006). Overexpression of p21 increases the resistance to apoptosis of alveolar epithelial cells (Inoshima et al., 2004), and the beneficial effects of lopinavir/ritonavir in VILI could represent the effects of the overexpression of this gene and the observed decrease in caspase-9. In contrast, absence of p21 was associated with more severe lung injury and increased numbers of apoptotic cells, as previously suggested (Yamasaki et al., 2008).

### The role of mechanical stretch

Mechanical overstretch could be an important pathogenic factor involved in p53/p21 activation. Several mechanisms have linked the mechanical load to a lung biological response, including oxidative stress and MAPK activation (Correa-Meyer et al., 2002). We focused on the role of the nuclear envelope as a critical structure regulating both mechanosensing and senescence. The mechanical load is transmitted from the extracellular matrix to the cytoskeleton and then to the nuclear membrane (Maurer and Lammerding, 2019). This causes a change in the nuclear lamina, reorganization of the underlying chromatin and DNA damage (Yang et al., 2017), either mediated by MAPK activation (Upadhyay et al., 2003) or by a direct mechanical effect (Nava et al., 2020). DNA damage is one of the triggers of the p53 pathway. In smooth muscle cells, stretch leads to p53 activation and upregulation of senescence markers (Mayr et al., 2002), resembling our findings. Similarly, HIV protease inhibitors, such as lopinavir/ritonavir, inhibit ZMPSTE-24, a protease responsible for Lamin-A maturation (Coffinier et al., 2007), preserving nuclear compliance, increasing p21 expression and decreasing stretch induced apoptosis and VILI (López-Alonso et al., 2018).

### Senescence in lung diseases

One of the known consequences of p53 activation is the cell switch towards a senescent phenotype. Lipopolysaccharide or bleomycin-induced lung injury increases the number of SA-β-galactosidase-positive cells and leads to cell cycle arrest (Sagiv et al., 2018). It has been shown that this activation has no detrimental effects in acute inflammation. However, blockade of the cell cycle has been associated with increased collagen deposition (Waters et al., 2018) and SASP may perpetuate lung inflammation (Kumar et al., 2014). Therefore, main features of abnormal lung repair after acute injury (limited cell proliferation, chronic inflammation and fibrosis) could be explained by a persistent senescent response. Inhibition of this response by selective deletion of Tp53 in Club cells ameliorated lung damage related to chronic inflammation (Sagiv et al., 2018), suggesting a novel mechanism amenable to treatment of lung diseases.

The acute nature of our model and its short-term lethality does not allow the identification of canonical senescence markers such as Senescence-associated β-galactosidase, as these require from days to weeks to be positive (Tominaga et al., 2019). However, we identified a set of early markers including changes in chromatin structure and gene expression. In a model of repair after VILI, lung *Cdkn1a* expression remained elevated up to two days of spontaneous breathing after injury (López-Alonso et al., 2018). Although it is unclear if these mechanisms may precipitate a full-blown senescent response in the long term, our results highlight the involvement of this molecular machinery in the early phase, and could be a therapeutic target to avoid late consequences.

### Clinical implications

Our findings have several implications regarding the pathogenesis of lung injury and its long-term consequences. First, the described p21 response may be beneficial in the acute phase, and could be pharmacologically manipulated using lopinavir. In a recent clinical trial in patients with lung disease caused by the SARS-CoV-2 coronavirus, lopinavir did not reduce mortality, but decreased the risk of ARDS development (Cao et al., 2020). However, the associated senescent response could worsen lung repair and long-term outcomes. Survivors after a prolonged ICU stay may have deleterious and prolonged sequels, including respiratory impairment (Heyland et al., 2005), neuropsychological disturbances (Girard et al., 2018) and muscle atrophy (Dos Santos et al., 2016), particularly the elderly (Heyland et al., 2015). The mechanisms responsible for these sequels are largely unknown and no effective therapies are currently available. Local activation of senescence and its paracrine/systemic spread could contribute to the pathogenesis of these sequels. As previously discussed, senescence may contribute to disordered lung repair. The confirmation of this framework could lead to the use of senolytics (Paez-Ribes et al., 2019, p.) in critically ill patients. However, due to the protective nature of senescence in the early phase (Chu et al., 2020), these treatments should be time-coordinated and modulated to optimize their effectiveness.

### Conclusions

We provide new evidence suggesting that acute lung damage activates p21 to limit apoptosis. This response appears to be trigged by the induction of DNA damage and linked to chromatin changes caused by mechanical overstretch. Interaction with the nuclear lamina may enhance this anti-apoptotic response. Although p21 activation may be beneficial in the acute phase of lung injury, the long-term effects must be taken into consideration as they could explain some of the long term sequels of critically ill patients.

## Experimental procedures

### Meta-analysis of transcriptomic data

To explore the main hypothesis, a pooled analysis of published transcriptomic data was performed, using a previously validated 55-gene expression signature of senescence (Hernandez-Segura et al., 2017) as main endpoint. Datasets reporting lung gene expression in animal models of acute lung injury and mechanical ventilation were obtained from public repositories (Gene Omnibus Expression - https://www.ncbi.nlm.nih.gov/geo/- and ArrayExpress - https://www.ebi.ac.uk/arrayexpress/-) using the following terms: “Stretch”, “Cyclic strain”, “Mechanical Ventilation”, “Lung”, and “Alveolar”. Fifty-one datasets were manually reviewed. Studies lacking a control group with intact, spontaneously breathing animals and those reporting less than 40 genes from the endpoint signature were excluded, so 11 datasets were finally used (Supplementary Table 1). When available, raw data was downloaded and normalized using the Robust Multiarray Average method (for Affymetrix microarrays) or normal-exponential background correction followed by quantile normalization (all the other platforms).

Normalized datasets were pooled using the Combat-Co-normalization using controls (COCONUT) algorithm (Sweeney et al., 2016). This method normalizes gene expression of the different datasets using an empirical Bayes fitting, but applied only to control samples (in this case, spontaneously breathing animals with intact lungs). Then the obtained normalization parameters are applied to the cases (i.e., those with lung injury)(Supplementary Figure 1). Three different signatures were studied, corresponding to 116 genes upregulated by p53, 14 genes downregulated by p21 (Fischer, 2017) and a set of 50 genes consistently up- and down-regulated in senescence (Hernandez-Segura et al., 2017). A meta-score was computed for each sample as the geometric mean of the upregulated genes minus the geometric mean of the downregulated genes in the signature. Meta-scores were finally compared among controls and animals with lung injury and/or mechanical ventilation.

### Animal models

Male, 12 week old C57Bl/6 mice, kept under pathogen-free conditions with free access to food and water, were used in all experiments. The Animal Research Committee of the Universidad de Oviedo evaluated and approved the study.

A two-hit lung injury model, based on chlorhydric acid instillation and mechanical ventilation, was studied. Animals were anesthetized with intraperitoneal ketamine and xylazine and orotracheally intubated using a 20G catheter, through which 50μL of chlorhydric acid (0.1N, pH=1.5) were instilled. Two hours after instillation, mice were randomly assigned to receive mechanical ventilation or not. Mice were ventilated with a pressure-controlled mode (peak inspiratory pressure 17 cmH_2_O, PEEP 2 cmH_2_O, respiratory rate 100 breaths/min) for 120 minutes.

Three additional series of experiments were performed. Mice lacking *Tp53* or *Cdkn1a* (p21, an endogenous inhibitor of cyclin-dependent kinases involved in the senescent response triggered by *Tp53*) and their wildtype littermates were subjected to the same model of injury, including acid instillation and mechanical ventilation. Genotypes were confirmed by PCR. In separate experiments, wildtype animals were treated with a single dose (200/50 mg/Kg) of lopinavir/ritonavir (a protease inhibitor that inhibits Zmpste24 and disrupts Lamin-A nuclear scaffolding, activating senescence pathways) or saline, administered intraperitoneally immediately after acid instillation, and then ventilated with the parameters described above.

### Tissue harvest

Mice were studied in three different conditions: baseline, 4 hours after chlorhydric acid instillation without mechanical ventilation and 4 hours after acid instillation including 2 hours of mechanical ventilation. Lungs were removed after exsanguination of anesthetized animals. A laparotomy was performed, the renal artery sectioned, the thorax opened and the heart-lungs removed *in bloc*. The left lung was instilled with 250 microliters of 4% phosphate-buffered paraformaldehyde, immersed in the same fixative for 24 hours, and then stored in 50% ethanol. The right lung was immediately frozen at −80°C for biochemical analyses.

### Patient samples

Paraffin-embedded lung tissue from autopsies of patients were obtained from the tissue bank at Hospital Universitario Central de Asturias, after signed consent from patients’ next of kin. ARDS was defined using the Kigali modification of the Berlin definition (Riviello et al., 2016), to include patients with lung injury but without mechanical ventilation and those without an arterial line. 13 samples were recovered (Supplementary Table 2).

### Histological studies

After fixation, tissues were embedded in paraffin and three slices with at least 1mm of separation between them were cut and stained with hematoxylin and eosin. A pathologist blinded to the experimental settings evaluated the degree and extension of lung damage using a predefined histological score (Blazquez-Prieto et al., 2015).

Additional lung sections were processed as previously described (Gonzalez-Lopez et al., 2011) for detection of myeloperoxidase- and Ki-67-positive cells, using specific antibodies (See Supplementary Table 3 for references). Images from three random fields (x200) were taken and then number of positive cells averaged.

For immunofluorescence studies, slides were deparaffinated and antigens retrieved in citrate buffer 0.1M (pH=9). The autofluorescence of the tissue was diminished using a Sudan black B solution and sections were permeabilized (0.1% Triton X-100 in PBS for 15 minutes), blocked (1% BSA in PBS) and incubated overnight at 4°C with the primary antibody (Supplementary Table 3). After 24 hours, the slices were incubated with the corresponding secondary fluorescent antibody at room temperature for 1 hour. Images were taken using a confocal microscopy (Leica SP8) at 400x and 630x. The number of positive and negative nuclei were automatically quantified using ImageJ software (NIH, USA).

Apoptotic cells in lung slices were detected by terminal deoxynucleotidyl transferase dUTP nick end labeling (TUNEL) as previously described (López-Alonso et al., 2018). Images from three random fields were acquired in a LEICA SP8 confocal microscope and the positive nuclei were counted and expressed as percentage of the total nuclei count.

### Western blot

Nuclei were extracted from fresh lung tissues and subsequently homogenized as described before (López-Alonso et al., 2018). The total amount of protein from nuclear extracts was quantified (BCA Protein Assay Kit, Pierce) and 15μg of each sample were loaded in SDS-polyacrylamide gels, electrophoresed at 120mV and electrotransferred onto PVDF membranes. After blockade with 5% non-fat dry milk, the membranes were incubated with primary antibodies against Caspase-9, Lamin-A/C, Lamin-B1, γH2AX, HP1α or H3 (Supplementary Table 3) in 3% non-fat dry milk overnight at 4°C. After 24 hours, the membranes were incubated with the corresponding peroxidase-conjugated secondary antibodies in 2.5% non-fat dry milk. Proteins were detected by chemiluminescence in a LAS-4000 Imaging system. The intensity of each protein band was quantified using ImageJ software (NIH, USA).

### Quantitative PCR

Lung fragments (2 mm x 2 mm) were homogenized with TRIZOL (Sigma, Poole, UK) and RNA precipitated by overnight incubation in isopropanol at −20°C. After 24 hours, samples were washed with ethanol and the RNA resuspended in RNAse-free water and quantified. One µg of total RNA was retrotranscribed into complementary cDNA using an RT-PCR kit (High capacity cDNA rt Kit, Applied Biosystems). Quantitative PCRs were carried out in triplicate of each sample using 40 ng of cDNA per well. Expression of *Plk3, Gdnf, Meis1, Il6, Tp53, Cdkn1a* (p21), *Cdkn2a* (p16), *Rb*, and *Gapdh* was quantified using Sybr-green Power up, (Fisher Scientific) and 10uM of the corresponding primers (Supplementary Table 4). The relative expression of each gene was calculated as 2^-^Δ^CT(gene of interest)–Δ CT(GAPDH)^.

### Statistical analysis

Data are shown as mean ± standard error of the mean. Differences between two groups were studied using a T test. Differences among more than two groups were assessed using an analysis of the variance (ANOVA). For SAHF counts, three slides per animal were counted (considered as technical replicates) and analyzed using a mixed-effects ANOVA. When significant, pairwise comparisons were done using the Tukey’s Honest Significant Difference test. A p-value lower than 0.05 was considered significant.

## Authors’ contributions

JBP, CHF, ILA and GMA designed the study. JBP, CHF, ILA, LAR, PMV, CLM, IC, CP and PJFM performed the experiments. GMA did the bioinformatic analysis. JBP, ILA, CHF, LAR, PJFM and GMA analyzed the results. CHF, ILA, LAR, PJFM, MS, JIS and GMA discussed the significance of the results. JIS, MS and GMA wrote the paper. GMA is the responsible of the integrity of the whole work.

