## Supplementary for "Activation of p21 limits acute lung injury and induces early senescence after acid aspiration and mechanical ventilation"

### Data supplement

**Supplementary table 1.** Datasets measuring gene expression in animal models of lung injury and mechanical ventilation. MV: Mechanical ventilation. DP: Driving pressure. VT: Tidal volume. LPS: Lipopolysaccharide.

| Dataset | Ref | Platform | Animal | Intervention | n |
|---|---|---|---|---|---|
| **GSE121550** | (López-Alonso et al. 2019) | GPL16570 | Mouse | Spontaneous breathing | 6 |
| | | | | MV (DP 15 $cmH_2O$), 2.5h | 6 |
| **GSE85269** | (López-Alonso et al. 2018) | GPL16750 | Mouse | Spontaneous breathing | 3 |
| | | | | MV (DP 15 $cmH_2O$), 2.5h | 3 |
| **GSE18341** | (Smith et al. 2010) | GPL1261 | Mouse | Spontaneous breathing | 8 |
| | | | | MV (VT15 ml/Kg), 2h | 7 |
| | | | | Inhaled LPS | 8 |
| | | | | MV (VT 15 ml/kg , 2h) + Inhaled LPS | 7 |
| **GSE11434** | (Wray et al. 2009) | GPL1261 | Mouse | Spontaneous breathing | 5 |
| | | | | MV (DP 20 $cmH_2O$), 3h | 5 |
| **GSE9368** | (Hong et al. 2008) | GPL1261 | Mouse | Spontaneous breathing | 3 |
| | | | | MV (VT 30 ml/kg), 4h | 3 |
| **GSE9314** | (Hong et al. 2008) | GPL1261 | Mouse | Spontaneous breathing | 4 |
| | | | | MV (VT 30 ml/kg), 4h | 4 |
| **GSE86229** | (Otulakowski et al. 2017) | GPL6246 | Mouse | Spontaneous breathing | 5 |
| | | | | MV (VT 35 ml/kg), 3h | 10 |
| **GSE31678** | (Park et al. 2012) | GPL1355 | Rat | Spontaneous breathing | 2 |
| | | | | MV (VT 18 ml/kg), 3h | 3 |
| **GSE7041** | (Nonas et al. 2007) | GPL1355 | Rat | Spontaneous breathing | 3 |
| | | | | MV (VT 20 ml/kg), 2h | 3 |
| **GSE9208** | (Papaiahgari et al. 2007) | GPL8321 | Mouse | Spontaneous breathing | 3 |
| | | | | MV (VT 30 ml/kg), 2h | 3 |
| **GSE2411** | (Altemeier et al. 2005) | GPL339 | Mouse | Spontaneous breathing | 6 |
| | | | | MV (VT 10 ml/kg), 4h | 6 |
| | | | | Inhaled LPS | 6 |
| | | | | MV (VT 10 ml/kg), 4h + Inhaled LPS | 6 |

**Supplementary Table 2.** Patients' characteristics. ARDS: Acute respiratory distress syndrome diagnosed according to the Kigali modification of the Berlin definition. MV: Mechanical ventilation.

| Age | Sex | ARDS | MV | Diagnoses |
|---|---|---|---|---|
| 55 | F | No | No | Brain lymphoma |
| 57 | M | No | No | Leriche syndrome. Cardiac arrest |
| 67 | M | No | No | Cardiac arrest |
| 68 | M | No | No | Pneumoconiosis |
| 64 | F | No | No | Infective endocarditis |
| 67 | F | Yes | No | Bone marrow aplasia. Nosocomial pneumonia |
| 68 | F | Yes | No | Community-acquired pneumonia. Septic shock |
| 48 | M | Yes | No | Liver cirrhosis. H1N1 influenza. Airway obstruction. |
| 59 | M | Yes | Yes | H1N1 influenza |
| 49 | M | Yes | Yes | H1N1 influenza |
| 53 | M | Yes | Yes | Nosocomial pneumonia |
| 42 | M | Yes | Yes | Community-acquired pneumonia |
| 68 | M | Yes | Yes | Community-acquired pneumonia |

**Supplementary Table 3.** Antibodies used in the study.

| | | |
|---|---|---|
| Western blotting | Primary antibodies | Anti-Caspase-9 (#9508, Cell signaling); dilution 1/1000 |
| | | Anti-Lamin A/C (sc-20681, Santa Cruz); dilution: 1/2000 |
| | | Anti-Lamin B1 (sc-374015, Santa Cruz); dilution: 1/1000 |
| | | Anti-YH2AX (sc-517348, Santa Cruz); dilution: 1/500 |
| | | Anti-HP1α (sc-130446, Santa Cruz); dilution: 1/500 |
| | | Anti-actin (sc-1616, Santa Cruz); dilution: 1/10000 |
| | | Anti-H3 (ab1791, Abcam); dilution: 1:10000 |
| | Secondary antibodies | Anti-rabbit IgG-HRP (sc-2004, Santa Cruz); dilution: 1/10000 |
| | | Anti-mouse IgG-HRP (sc-2005, Santa Cruz); dilution: 1/10000 |
| Immuno-histo-chemistry | Primary antibodies | Anti-myeloperoxidase (A0398, Dako) |
| | | Anti-Ki67 (GA626, Dako) |
| | Secondary antibodies | Anti-mouse IgG-HRP (P044701-2, Dako) |
| | | Anti-rabbit IgG-HRP (P044801-2, Dako) |
| Immuno-fluorescence | Primary antibodies | Anti-macro-H2A (15825913, Fisher); dilution: 1/250 |
| | | Anti-Lamin A/C (sc-20681, Santa Cruz); dilution: 1/250 |
| | Secondary antibodies | Anti-rabbit IgG-Alexa Fluor 594 (A-21207, Invitrogen); dilution:1/2000 |
| | | Anti-mouse IgG-FITC (sc-2010, Santa Cruz); dilution: 1/500 |

**Supplementary Table 4.** Primers used for quantitative PCR (FW: Forward; RV: Reverse).

| Gene | Species | Direction | Sequence |
|---|---|---|---|
| ***Tp53*** | Mouse | FW | 5’-CTCTCCCCCGCAAAAGAAAAA-3’ |
| | | RV | 5’-CGGAACATCTCGAAGCGTTTA-3’ |
| ***Cdkn1a*** | Mouse | FW | 5’-GGAACATCTCAGGGCCGAAA-3’ |
| | | RV | 5’-AAGACCAATCTGCGCTTGGA-3’ |
| ***Cdkn2a*** | Mouse | FW | 5’-CGCAGGTTCTTGGTCACTGT-3’ |
| | | RV | 5’-TGTTCACGAAAGCCAGAGCG-3’ |
| ***Il6*** | Mouse | FW | 5′-ACCACTTCACAAGTCGGAGG-3′ |
| | | RV | 5’-TGCAAGTGCATCATCGTTGT-3’ |
| ***Plk3*** | Mouse | FW | 5’-GCGCGAGAAGATCCTAAATG-3’ |
| | | RV | 5’-CTCTGGTTCCAACAGGGTGT-3’ |
| ***Gdnf*** | Mouse | FW | 5’-GACTTGGGTTTGGGCTATGA-3’ |
| | | RV | 5’-AACATGCCTGGCCTACTTTG-3’ |
| ***Meis1*** | Mouse | FW | 5’-AAGGTGATGGCTTGGACAAC-3’ |
| | | RV | 5’-TGTGCCAACTGCTTTTTCTG-3’ |
| ***Rb1*** | Mouse | FW | 5’-GCAGTCCAAGGATGGAGAAG-3’ |
| | | RV | 5’-ACAGGGCAAGGGAGGTAGAT-3’ |
| ***Gapdh*** | Mouse | FW | 5’-GTGCAGTGCCAGCCTCGTCC-3’ |
| | | RV | 5’-GCCACTGCAAATGGCAGCCC-3’ |
| ***TP53*** | Human | FW | 5’-GTGCAGCTGTGGGTTGATTC-3’ |
| | | RV | 5’-ACCATCGCTATCTGAGCAGC-3’ |
| ***CDKN1A*** | Human | FW | 5’-TGTCCGTCAGAACCCATGC-3’ |
| | | RV | 5’-AAAGTCGAAGTTCCATCGCTC-3’ |
| ***GAPDH*** | Human | FW | 5’-TCGGAGTCAACGGATTTGGTCGT-3’ |
| | | RV | 5’-TGCCATGGGTGGAATCATATTGGA-3’ |

**Supplementary figure 1.** Gene expression histograms before (single-study normalization) and after normalization with the COCONUT algorithm (conjoint normalization). Controls were spontaneously breathing animals with healthy lungs. Cases were all those with acute lung injury.

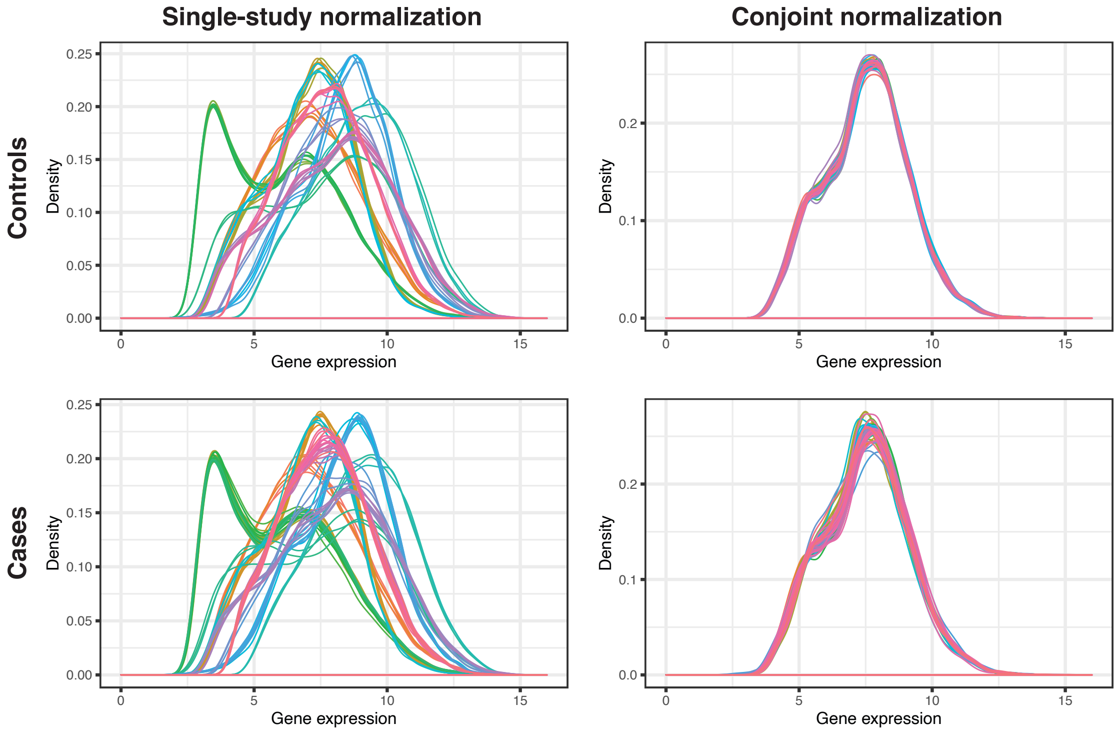

**Supplementary figure 2.** Heatmaps showing pooled expression of genes included in specific signatures regulated by p53 (A), p21 (B) or up- and down-regulated by senescence (C, top and bottom panels respectively).

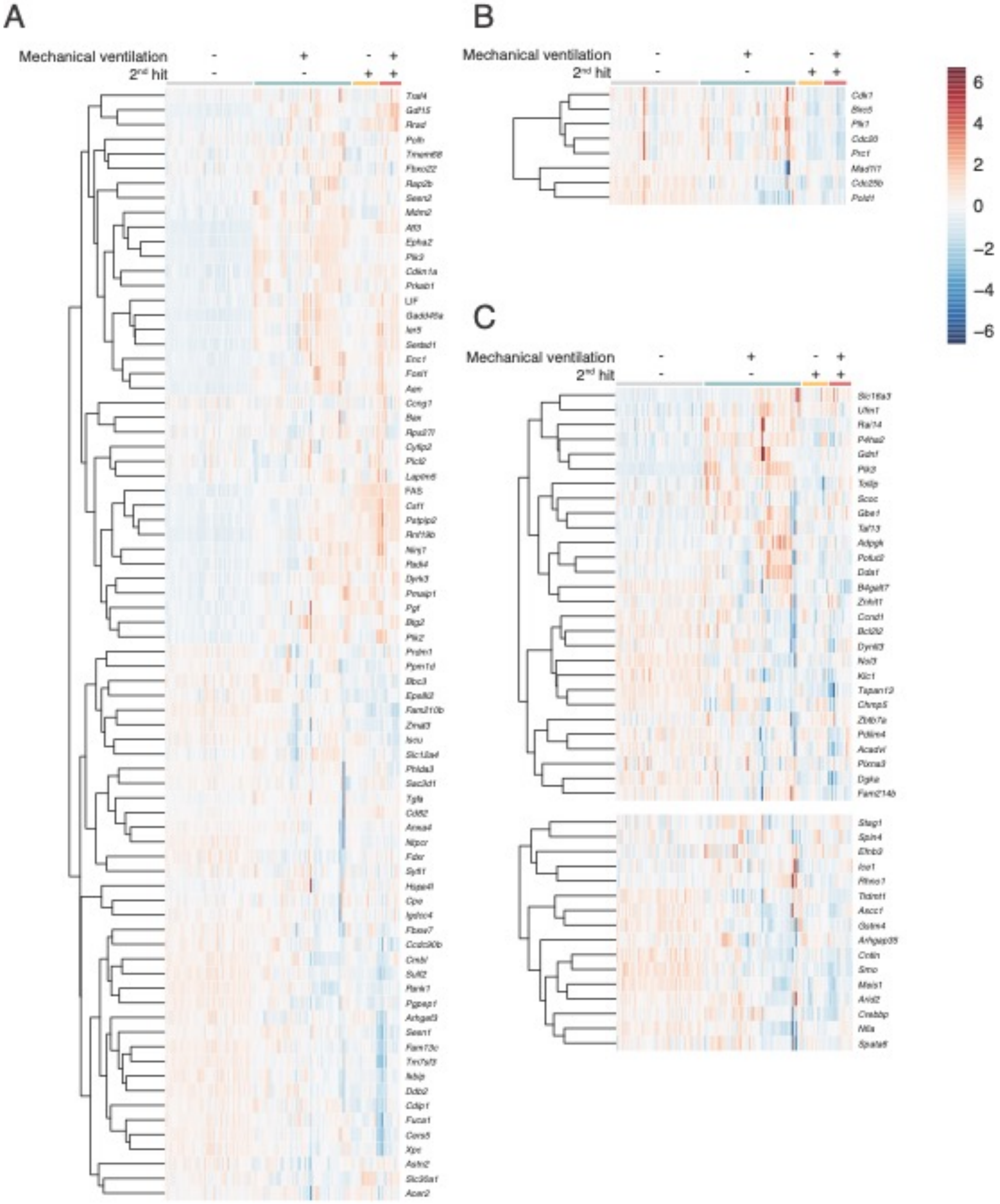

**Supplementary figure 3.** Representative images of histological slides showing MPO (A), Ki-67 (B) and TUNEL (C) positive cells.

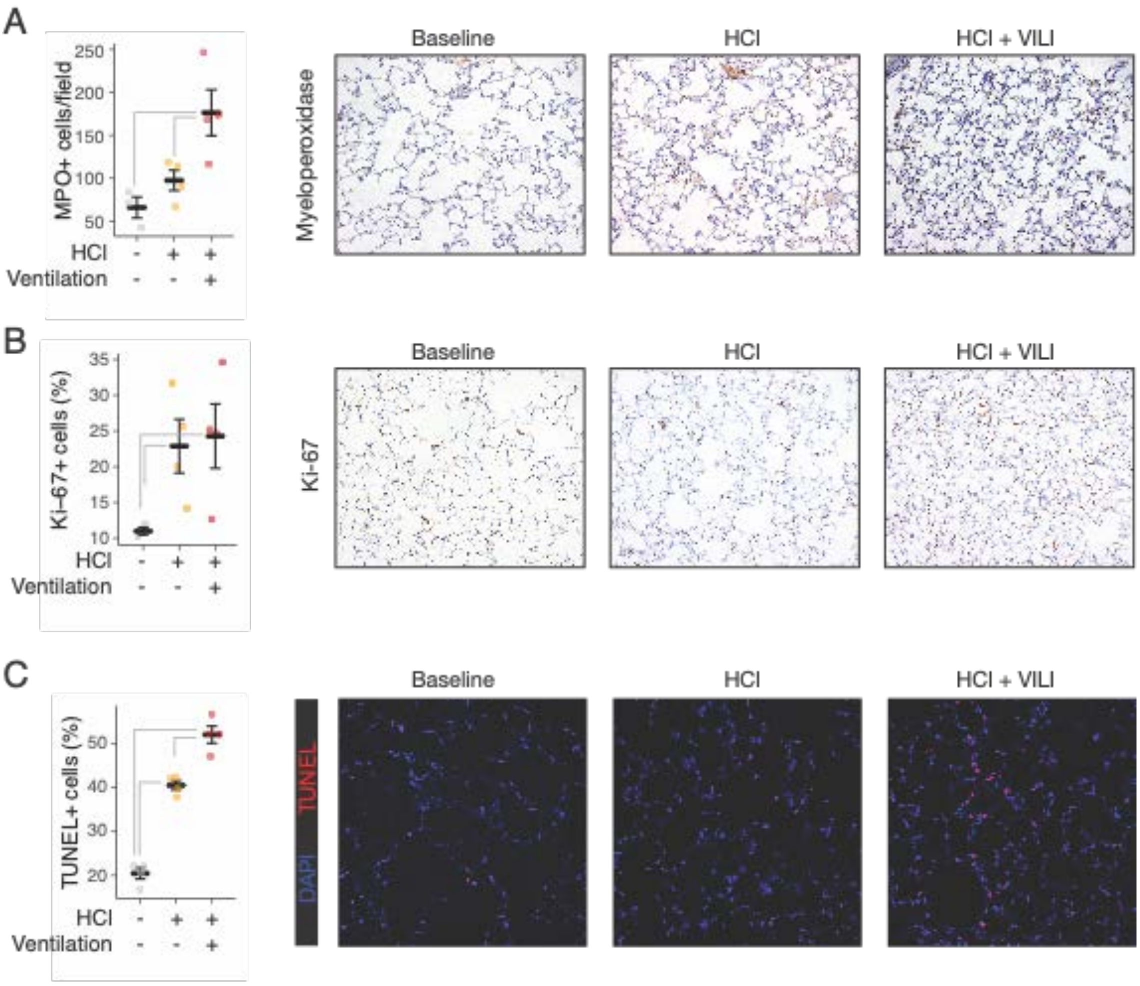

**Supplementary figure 4.** Representative histological sections of *Tp53*$^{+/+}$ and *Tp53*$^{-/-}$ mice after acid instillation and mechanical ventilation. There were no significant differences in the severity of lung injury between genotypes.

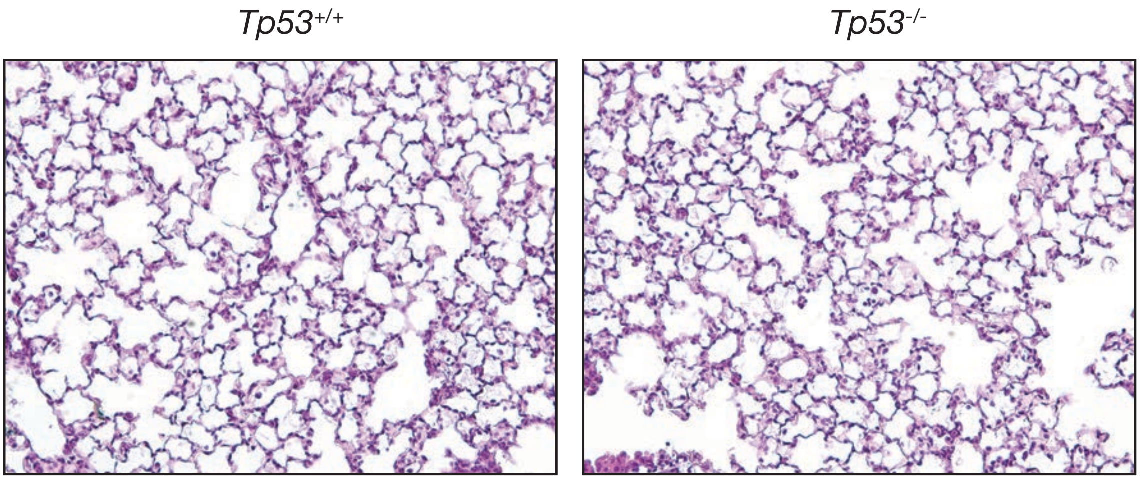
